# Trehalose recycling promotes energy-efficient mycomembrane reorganization in nutrient-limited mycobacteria

**DOI:** 10.1101/758177

**Authors:** Amol Arunrao Pohane, Caleb R. Carr, Jaishree Garhyan, Benjamin M. Swarts, M. Sloan Siegrist

**Affiliations:** Department of Microbiology, University of Massachusetts, Amherst, Massachusetts, USA; Department of Chemistry and Biochemistry, Central Michigan University, Mount Pleasant, Michigan, USA; Molecular and Cellular Biology Graduate Program, University of Massachusetts, Amherst, Massachusetts, USA

## Abstract

The mycomembrane layer of the mycobacterial cell envelope is a barrier to environmental, immune and antibiotic insults. We find that there is mycomembrane remodeling along the periphery of nutrient-starved, non-replicating mycobacterial cells. Remodeling is supported by recycling of trehalose, a non-mammalian disaccharide that shuttles long-chain mycolate lipids to the mycomembrane. In the absence of trehalose recycling, mycomembrane synthesis continues but mycobacteria experience ATP depletion, enhanced respiration and redox stress. Redox stress from depletion of the trehalose pool is suppressed in a mutant that lacks the OtsAB *de novo* trehalose synthesis pathway. Our data suggest that trehalose recycling alleviates the energetic burden of mycomembrane remodeling. Loss of trehalose salvage is known to attenuate *M. tuberculosis* during infection and render the bacterium more susceptible to a variety of drugs. Recycling pathways are emerging targets for sensitizing resource-limited bacterial pathogens to host and antibiotic stress.

## Introduction

The mycobacterial cell envelope is comprised of covalently-bound peptidoglycan, arabinogalactan and mycolic acids, as well as intercalated glycolipids and a thick capsule (Puffal et al., 2018). The mycolic acids attached to the arabinogalactan and the noncovalent glycolipids respectively form the inner and outer leaflets of the mycomembrane, a distinctive outer membrane present in members of the *Corynebacterineae* suborder. The mycomembrane is a key determinant of envelope permeability and home to a variety of immunomodulatory lipids and glycolipids (Gebhardt et al., 2007; Lee et al., 2016; Yang et al., 2014). There is substantial evidence that the mycomembrane is remodeled *in vivo* and in response to host-mimicking stresses, conditions in which mycobacterial growth and envelope synthesis are presumed to be slow or nonexistent (Bacon et al., 2014; Baker and Abramovitch, 2018; Bhamidi et al., 2011; Eoh et al., 2017; Galagan et al., 2013; Kieser and Rubin, 2014; Ojha et al., 2010; Shui et al., 2007; Yang et al., 2014). While these studies have elucidated bulk changes in mycomembrane composition, the dynamics and subcellular distribution of the molecular transitions have not been characterized. It is also unclear in most cases whether the alterations are solely catabolic, or whether anabolic reactions also contribute to changes in mycomembrane composition under stress.

Recycling pathways are likely to be at the nexus of stress-triggered mycomembrane reorganization. Mycolic acids are ligated to the non-mammalian disaccharide trehalose in the cytoplasm (Quemard, 2016). Once transported to the periplasm, trehalose monomycolate (TMM) donates its mycolic acid to arabinogalactan, forming arabinogalactan mycolates (AGM), or to an acceptor TMM, forming trehalose dimycolate (TDM; **Figure 1A**). Both processes release free trehalose. TDM can also be degraded by TDM hydrolase (TDMH) into TMM and free mycolic acids, the latter of which are an important component of biofilm extracellular matrix in mycobacteria (Holmes et al., 2019; Ojha et al., 2010). While a salvage mechanism for mycolic acids is still under debate (Cantrell et al., 2013; Dunphy et al., 2010; Forrellad et al., 2014; Wilburn et al., 2018), recapture of trehalose occurs via the LpqY-SugABC transporter (Kalscheuer et al., 2010b). Depending on the specific environmental demand, mycobacteria may funnel reclaimed trehalose back to central carbon metabolism, to generate intermediates for glycolysis or the pentose phosphate pathway, or store it in the cytoplasm, possibly as a stress protectant or compatible solute (Eoh et al., 2017; Lee et al., 2019; Nobre et al., 2014; Shleeva et al., 2017). An additional but unexplored potential fate for recaptured trehalose is direct reincorporation into TMM or other glycoconjugates destined for the cell surface. Thus, trehalose is a key molecule that connects extracellular processes—mycomembrane synthesis, remodeling and degradation—to the metabolic status of the mycobacterial cell.

**Figure 1.**
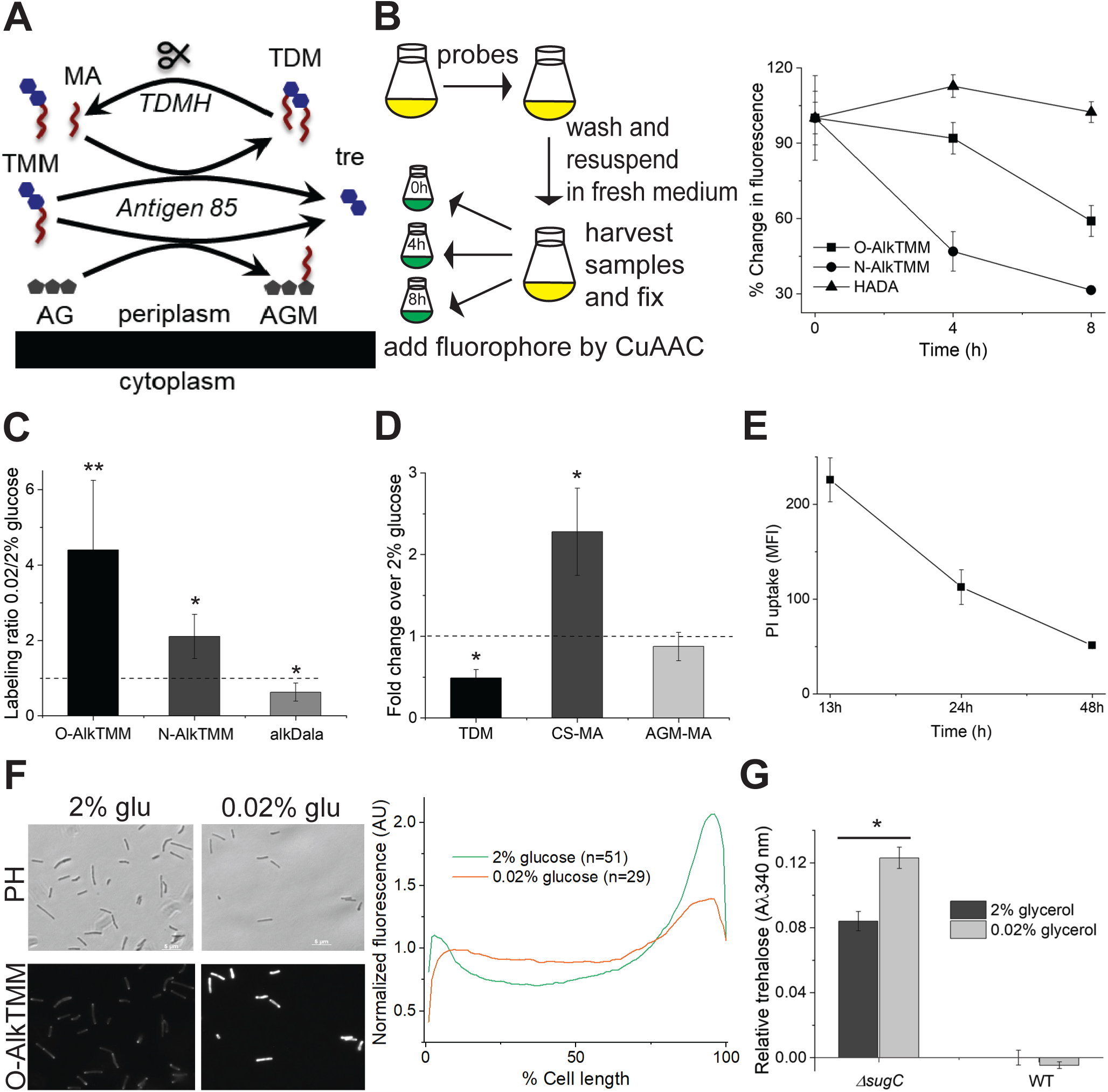
Mycomembrane synthesis and degradation are active under carbon limitation. (A) Cartoon of mycomembrane synthesis and degradation. TMM, trehalose monomycolate; TDM, trehalose dimycolate; AG, arabinogalactan; AGM, arabinogalactan mycolates; MA, free mycolic acids. (B) TDM turnover under nutrient deprivation. *M. smegmatis* cultured in 0.02% glucose-supplemented medium was labeled with metabolic probes O-AlkTMM (primarily labels AGM), N-AlkTMM (labels TDM) or HADA (labels cell wall peptidoglycan), then washed. Aliquots were taken at the indicated time points and fixed with 2% formaldehyde. Alkynes were detected by copper-catalyzed azide-alkyne cycloaddition (CuAAC) reaction with carboxyrhodamine 110 azide. Fluorescence was quantitated by flow cytometry, with the median fluorescence intensities normalized to the initial, 0 h time point for each probe. Experiment was performed three times in triplicate; one representative experiment shown. (C) Metabolic labeling of *M. smegmatis* in 0.02% glucose-supplemented medium with O-AlkTMM, N-AlkTMM and alkDala (labels peptidoglycan). Alkynes were detected by CuAAC reaction with carboxyrhodamine 110 azide. Data were normalized to labeling in 2% glucose-supplemented medium and plotted from four independent experiments. (D) Quantitation of thin-layer chromatography (TLC) of different mycomembrane components for *M. smegmatis* in 0.02%-supplemented medium. TDM, trehalose dimycolate; CS-MA, free, culture supernatant mycolic acids; AGM-MA, mycolic acids released from arabinogalactan. TLCs were scanned and processed in ImageJ (Schneider et al., 2012). Data are normalized to TLCs from samples taken from *M. smegmatis* cultured in 2% glucose-supplemented medium and plotted from three independent experiments. Representative TLC, **Figure S2**. (E) Propidium iodide staining of *M. smegmatis* upon adaptation to 0.02% glucose-supplemented medium. Fluorescence was quantitated by flow cytometry and median fluorescence intensity (MFI) was plotted. Experiment was performed three times in triplicate; one representative experiment shown. (F) O-AlkTMM labeling of *M. smegmatis* AGM in 2% or 0.02% glucose-supplemented medium. Alkynes were detected by CuAAC reaction with carboxyrhodamine 110 azide. Left, fluorescence microscopy. Scale bars, 5 μM. Right, cellular fluorescence was quantitated for cells lacking visible septa. Signal was normalized to both cell length and to total fluorescence intensity. Cells were oriented such that the brighter pole is on the right hand side of the graph. Experiment was performed three times; one representative experiment shown. (G) Quantification of trehalose from supernatants of wild-type and *ΔsugC M. smegmatis* cultured in 2% or 0.02% glycerol supplemented medium. Experiment was performed at least three times in triplicate; one representative experiment shown. Error bars, standard deviation. Statistical significance of 0.02% vs. 2% glucose or glycerol samples was assessed by two-tailed Student’s t test. *, p<0.05; **, p<0.005.

Here we show that mycomembrane remodeling triggered by nutrient limitation comprises both synthesis and degradation of AGM and TDM. While remodeling continues in the absence of trehalose recycling, compensatory *de novo* synthesis of trehalose via the OtsAB pathway upsets the redox and metabolic balance of the cell, phenotypes similar to those induced by futile cycling (Adolfsen and Brynildsen, 2015; Brynildsen et al., 2013) or some bactericidal antibiotics (Yang et al., 2017). Loss of trehalose recycling sensitizes *M. tuberculosis* and *M. smegmatis* to multiple drugs (Danelishvili et al., 2017) and attenuates *M. tuberculosis in vivo* (Kalscheuer et al., 2010b). Given the metabolic and energetic costs associated with synthesizing the cell envelope from scratch, salvage pathways may be particularly important for bacterial resilience to stress.

## Results

### Mycomembrane synthesis and degradation are active under carbon limitation

Decreased TDM abundance has been reported for mycobacteria growing in biofilms or adapting to hypoxia or nutrient limitation (Galagan et al., 2013; Lee et al., 2019; Ojha et al., 2010; Yang et al., 2014). As uncontrolled TDM hydrolysis results in cell lysis (Ojha et al., 2010; Yang et al., 2013), we sought to understand the kinetics of TDM turnover under stress. TMM donates mycolic acids to other molecules of TMM, to form the TDM glycolipid, or to arabinogalactan, to form covalent arabinogalactan mycolates (AGM). The TMM-mimicking probe N-AlkTMM specifically incorporates into TDM because the amide linkage permits mycolic acid acceptance but not donation of the alkyne-appended lipid chain (Foley et al., 2016). To track TDM hydrolysis under carbon limitation we performed a pulse chase experiment in which we labeled *M. smegmatis* with N-AlkTMM for 12 hours in low (0.02%) glucose-supplemented 7H9 medium, washed, then transferred to 7H9 lacking both the probe and glucose (**Figure 1B,** left). Alkyne-labeled TDM was detected on fixed cells at 0, 4 and 8 hours post-transfer by copper-catalyzed azide-alkyne cycloaddition (CuAAC) with a fluorescent azide label. We found that TDM labeling decreased by about 3-fold in this time period (**Figure 1B**). Fluorescence derived from D-amino acid-labeled cell wall peptidoglycan remained steady, however, consistent with limited bacterial growth under this condition (**Figures 1B, S1C**).

Under acid stress, non-replicating but metabolically-active *M. tuberculosis* make new TDM (Baker and Abramovitch, 2018). We found that N-AlkTMM uptake (no chase) increased approximately two-fold in low glucose medium (**Figure 1C**). However, a decline in the steady-state abundance of the molecule (**Figures 1D, S2A**) suggested that enhanced synthesis is outweighed by the TDM turnover observed in the pulse-chase experiment (**Figure 1B**).

We wondered whether there might be additional changes in mycomembrane metabolism. O-AlkTMM is also a TMM-mimicking probe but features an ester-linked lipid chain. While the molecule can serve as either an alkyne-lipid donor or acceptor, ∼90% of labeling from this probe is present in the *M. smegmatis* AGM cellular fraction (Foley et al., 2016). O-AlkTMM uptake was enhanced in low glucose medium to a greater extent than N-AlkTMM (**Figure 1C**). The fluorescence signal derived from this probe was also more persistent than N-AlkTMM in a no probe, no glucose chase (**Figure 1B**).

A variety of carbohydrates can serve as mycolate acceptors, including glucose (Gavalda et al., 2014; Matsunaga et al., 2008). High levels of glucose in the growth medium might therefore suppress O-AlkTMM labeling of the cell surface by competing with arabinogalactan. While *M. smegmatis* grew similarly in 7H9 with high (2%) vs. medium (0.2%) glucose supplementation, O-AlkTMM labeling in the former condition was lower. However, O-AlkTMM labeling was similar for *M. smegmatis* in 0.2% or 0.02% glucose or acetate, despite sluggish bacterial replication under the low carbon conditions (**Figures S1A, S1C**). Thus, incorporation of O-AlkTMM into AGM is suppressed in high glucose, likely because the alkyne-fatty acid from the probe is transferred to the unanchored glucose and washed away. Nonetheless our data indicate that substantial AGM synthesis occurs in growth-limiting amounts of glucose or acetate. As the steady-state abundance of the molecule did not change in carbon-limited medium (**Figure 1D, S2C**), these experiments also suggest that AGM synthesis is balanced by the turnover that we observed by pulse-chase (**Figure 1B**).

We previously showed that the fluorescent D-amino acid HADA and alkDala incorporate into *M. smegmatis* peptidoglycan via both cytoplasmic and L,D-transpeptidase enzymes (Garcia-Heredia et al., 2018). HADA and alkDala labeling roughly correlated with mycobacterial growth rate under different amounts of glucose or acetate (**Figures 1C, S1B, S1C**). Suppressed levels of peptidoglycan synthesis or remodeling during carbon limitation stood in contrast to active mycomembrane metabolism.

### AGM synthesis occurs along the periphery of the mycobacterial cell during carbon limitation

TDM hydrolysis enhances envelope permeability in oleic acid- and glucose-deprived *M. tuberculosis* (Yang et al., 2014). Surprisingly, despite an analogous decrease in TDM abundance (**Figure 1D**), *M. smegmatis* became less permeable to propidium iodide when cultured in glucose-limited medium (**Figure 1E**). Global AGM levels have also been linked to mycobacterial permeability (Jackson et al., 1999). While AGM abundance was relatively unaffected in glucose-deprived medium (**Figures 1D, S2C**), our data suggest that the apparent stasis belies active synthesis and degradation (**Figures 1B, 1C**). We wondered whether AGM remodeling might impact its spatial distribution, which in turn could alter cell permeability.

Mycobacteria growing in nutrient-replete medium construct their cell envelope in gradients that emanate from the poles and continue along the sidewall (Foley et al., 2016; Garcia-Heredia et al., 2018). While polar peptidoglycan synthesis promotes cell elongation, sidewall synthesis occurs in response to cell wall damage. We hypothesized that the AGM synthesis that we observe under carbon deprivation (**Figure 1C**) is a cell-wide response, similar to peptidoglycan repair. Quantitative fluorescence microscopy revealed that O-AlkTMM labeling of *M. smegmatis* growing in carbon-replete medium comprised polar gradients (**Figure 1F**) as expected (Foley et al., 2016; Garcia-Heredia et al., 2018). However, in slow or non-growing, carbon-deprived *M. smegmatis*, O-AlkTMM-labeled species were more evenly distributed around the periphery of the cell. This observation suggests that AGM synthesis fortifies the mycomembrane along the sidewall as mycobacteria adapt to carbon deprivation.

### Trehalose cycling supports mycomembrane metabolism during carbon starvation

Mycomembrane synthesis centers on the mycolic acid donor trehalose monomycolate (TMM). Prior to its export to the periplasm, TMM is synthesized in the cytoplasm by the ligation of a mycolic acid to trehalose (Kalscheuer and Koliwer-Brandl, 2014). *De novo* synthesis of mycolic acids and trehalose is both energy- and resource-intensive; recycling pathways for both molecules have been shown or proposed (Cantrell et al., 2013; Forrellad et al., 2014; Kalscheuer et al., 2010b). We hypothesized that nutrient-starved mycobacteria might buffer the costs of TMM synthesis by enlisting such pathways. As the recycling mechanism for mycolic acids is still controversial (Dunphy et al., 2010; Wilburn et al., 2018), we focused on the role of trehalose uptake.

Trehalose released as a byproduct of extracellular mycomembrane metabolism is recycled via the LpqY-SugABC transporter (Kalscheuer et al., 2010b). At least two different processes liberate trehalose: ligation of mycolic acids from TMM to arabinogalactan to form AGM and transfer of mycolic acids from TMM to another molecule of TMM to form TDM. Breakdown of TDM by the TDM hydrolase (TDMH) yields TMM and mycolic acids (Holmes et al., 2019; Ojha et al., 2010; Yang et al., 2013), so subsequent use of TMM in the forgoing reactions would also release trehalose. Our metabolic labeling results suggested that all of these processes are active as *M. smegmatis* adapts to carbon limitation (**Figure 1**). We were unable to measure extracellular trehalose levels in wild-type *M. smegmatis*, presumably because LpqY-SugABC rapidly internalizes the disaccharide (Kalscheuer et al., 2010b). However by using *ΔsugC M. smegmatis*, a strain that lacks a functional trehalose transporter, we were able to detect elevated levels of trehalose in the supernatant when bacteria were grown in carbon-limited conditions (**Figure 1G**; note that we used glycerol as the carbon source as glucose interferes with the assay). We also found that free mycolic acids accumulated in the supernatant of low glucose cultures, as expected from TDM degradation (**Figures 1D, S2B**). Together our data indicate that trehalose is liberated upon reorganization of the mycomembrane (**Figure 1A**).

After being transported by LpqY-SugABC (Kalscheuer et al., 2010b) and metabolized by trehalase (Nobre et al., 2014) or TreS (Eoh et al., 2017; Kalscheuer and Koliwer-Brandl, 2014; Kalscheuer et al., 2010a; Miah et al., 2013) **(Figure 2A**), high concentrations of exogenously-supplied trehalose can support mycobacterial growth in the absence of other carbon sources (Kalscheuer et al., 2010b). We recovered similar colony-forming units (CFU) for *ΔsugC*, *Δtre*, *ΔtreS* and wild-type *M. smegmatis* from 1, 2, 4 or 6 days in low glucose (**Figures 2B, 2C**). These data suggest that trehalose catabolism is not required for viability under carbon deprivation. Given that both the optical density and colony-forming units of wild-type *M. smegmatis* remain steady (**Figures 2B, 2C, S1C**), trehalose recovered from the mycomembrane also does not fuel appreciable cell growth under this condition.

**Figure 2.**
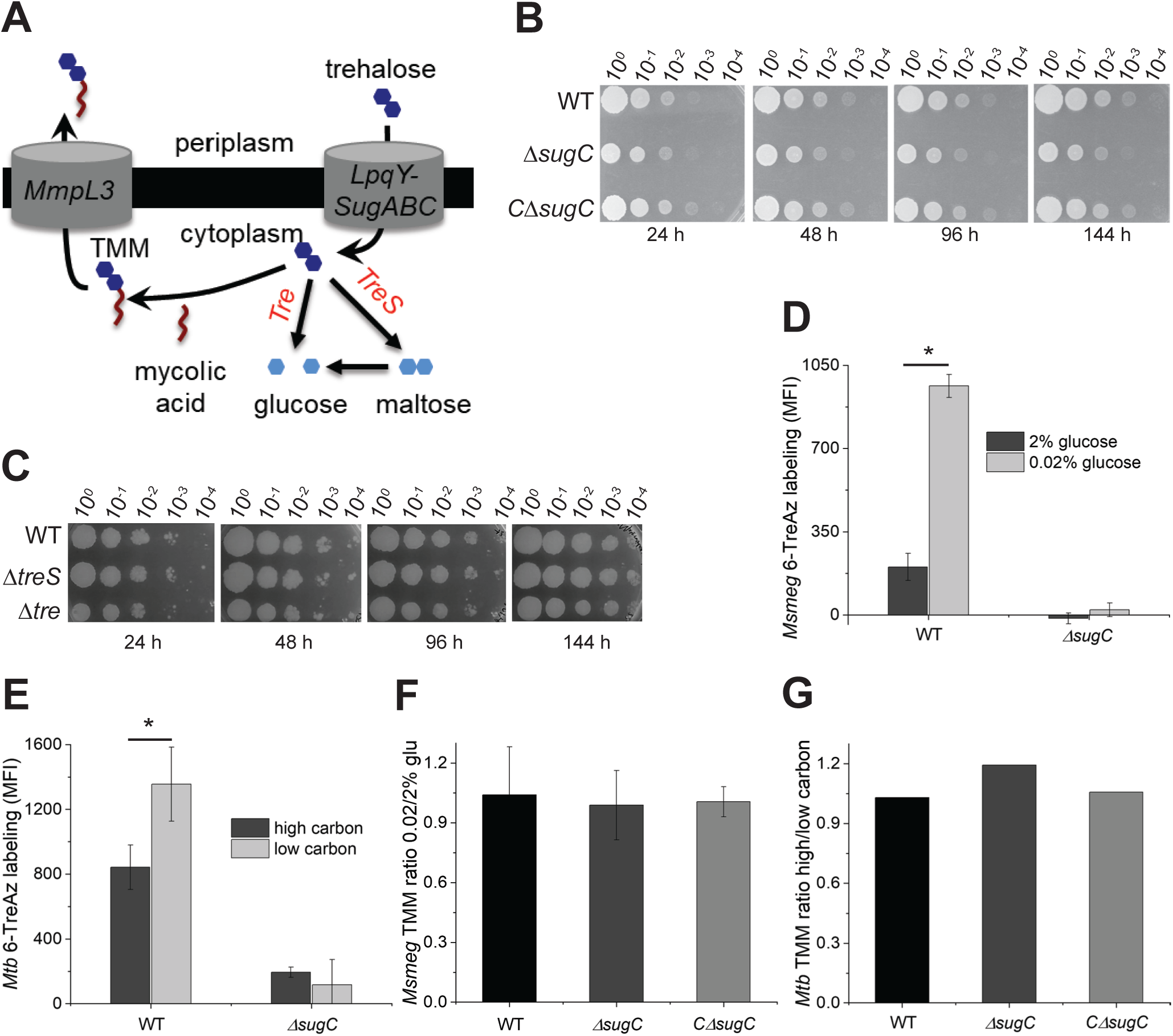
Trehalose cycling supports mycomembrane metabolism during carbon starvation. (A) Cartoon showing the utilization of recycled trehalose either by catabolic pathways mediated by trehalase (Tre) or TreS or in the synthesis of trehalose monomycoate (TMM). (B) and (C), Survival of wild-type, *ΔsugC,* complemented *ΔsugC* (C*ΔsugC*), *ΔtreS* and *Δtre M. smegmatis* in 0.02% glucose-supplemented medium. 10-fold serial dilutions were plated at the indicated time points. Experiment was performed two times with similar results; one experiment shown. (D) and (E), 6-TreAz labeling of wild-type and *ΔsugC M. smegmatis* (*Msmeg*) and *M. tuberculosis* (*Mtb*) cultured in low or high carbon medium. Azides were detected by strain-promoted azide-alkyne cycloaddition (SPAAC) with DBCO-Cy5 label. Fluorescence was detected by flow cytometry, with median fluorescence intensity (MFI) values from controls lacking 6-TreAz (but subjected to SPAAC) subtracted from sample MFI. Experiment was performed at least three times in triplicate; one representative experiment shown. (F) and (G), comparison of TMM abundance in *M. smegmatis* and *M. tuberculosis* cultured in grown low and high carbon medium. TLCs were scanned and processed in ImageJ (Schneider et al., 2012). Data are normalized to TLCs from mycobacteria cultured in high carbon medium and plotted from two (*M. tuberculosis*) or three (*M. smegmatis*) independent experiments. Representative TLCs, **Figure S3**. Error bars, standard deviation. Statistical significance of low vs. high carbon samples was assessed by two-tailed Student’s t test. *, p<0.05.

In hypoxic and biofilm cultures of *M. tuberculosis*, TMM and TDM levels decrease (Eoh et al., 2017; Galagan et al., 2013; Lee et al., 2019). Glycolipid turnover occurs rapidly in the former, within 4 hours (Eoh et al., 2017), and slowly in the latter, within 16 days (Lee et al., 2019). We did not observe a net decrease in TMM under our conditions (**Figures 2F, 2G**) despite an increase in TMM-consuming AGM and TDM remodeling (**Figure 1C**). We posited that TMM pools might be replenished by recycled trehalose. Metabolic incorporation of exogenous 6-azido-trehalose (6-TreAz) by *M. smegmatis* or *M. bovis* BCG requires uptake by LpqY-SugABC (Swarts et al., 2012) (**Figure S3A**). We found that 6-TreAz labeling was enhanced in slow-growing, glucose-starved *M. smegmatis* (**Figure 2D**) or oleic acid- and glucose-starved *M. tuberculosis* (**Figure 2E**, (Yang et al., 2014)). As incorporation of the metabolite was respectively abolished or diminished in *ΔsugC M. smegmatis* (**Figure 2D**, (Swarts et al., 2012)) or *M. tuberculosis* (**Figure 2E**), enhanced 6-TreAz labeling under carbon limitation indicates an increase in trehalose recycling.

6-TreAz recovered by the LpqY-SugABC transporter may remain intact in the cytoplasm or be converted to azido-TMM and transported outside of the cell (**Figure 2A**, (Swarts et al., 2012). Although it has not been reported, it is possible that the probe incorporates into other trehalose-bearing molecules in the mycobacterial envelope (Nobre et al., 2014). To tune our detection for the cell surface, we selected DBCO-Cy5 as the fluorescent, azide-reactive label because the localized charge on the sulfonated cyanine dye confers poor membrane permeability (Yang and Hinner, 2015). Thus, the enhanced 6-TreAz labeling that we observed for *M. smegmatis* and *M. tuberculosis* during carbon limitation (**Figures 2D, 2E**) strongly suggests that at least some of the recycled trehalose is converted into an envelope component(s). Given that 1) TMM and TDM are the only known trehalose-containing glycoconjugates shared by both *M. smegmatis* and *M. tuberculosis*, and that 2) TDM is not labeled by 6-TreAz (Swarts et al., 2012), we conclude that TMM is the most likely target. As steady-state TMM levels remained relatively constant in both species (**Figures 2F, 2G, S3B, S3C**), enhanced conversion of 6-TreAz to azido-TMM further suggests that trehalose recycling under carbon deprivation helps to maintain TMM levels. Taken together, our data support a model in which trehalose cycles in and out of trehalose mycolates and inside and outside of the cell to remodel the mycomembrane in carbon-deprived mycobacteria.

### Mycomembrane reorganization under carbon deprivation can occur in the absence of trehalose cycling

Our experiments suggest that trehalose cycling contributes to mycomembrane reorganization during carbon limitation. However, loss of trehalose import by LpqY-SugABC did not impact the abundance of TMM, TDM or AGM (**Figures S2A, S2B, S2C, S4A, S4B**); synthesis of AGM or TDM (**Figure S4C**); degradation of TDM (**Figure S4D**); permeability (**Figure S4E**). The absence of measurable changes in mycomembrane metabolism or composition were consistent with earlier work showing that *ΔsugC* and *ΔlpqY M. tuberculosis* do not have detectable changes in the glycolipid composition of their mycomembrane (Kalscheuer et al., 2010b). These data show that mycomembrane reorganization can occur in the absence of trehalose recycling.

### Trehalose recycling promotes *M. tuberculosis* survival in macrophages

While trehalose recycling was dispensable for mycomembrane remodeling and survival in an *in vitro* model of carbon limitation (**Figures S4**, **2B**), we hypothesized that it might be important for withstanding other stressors. Deletion of *sugC* or *lpqY* inhibits *M. tuberculosis* replication in the acute phase of murine infection (Kalscheuer et al., 2010b). Transposon insertions in *sugABC or lpqY* also attenuate pooled *M. tuberculosis* growth in IFN-γ-activated and unactivated C57BL/6 bone marrow-derived macrophages (Rengarajan et al., 2005). We confirmed that *ΔsugC M. tuberculosis* was defective for growing in these host cells (immortalized) and that this phenotype was reversed by genetic complementation (**Figure 3A**).

**Figure 3.**
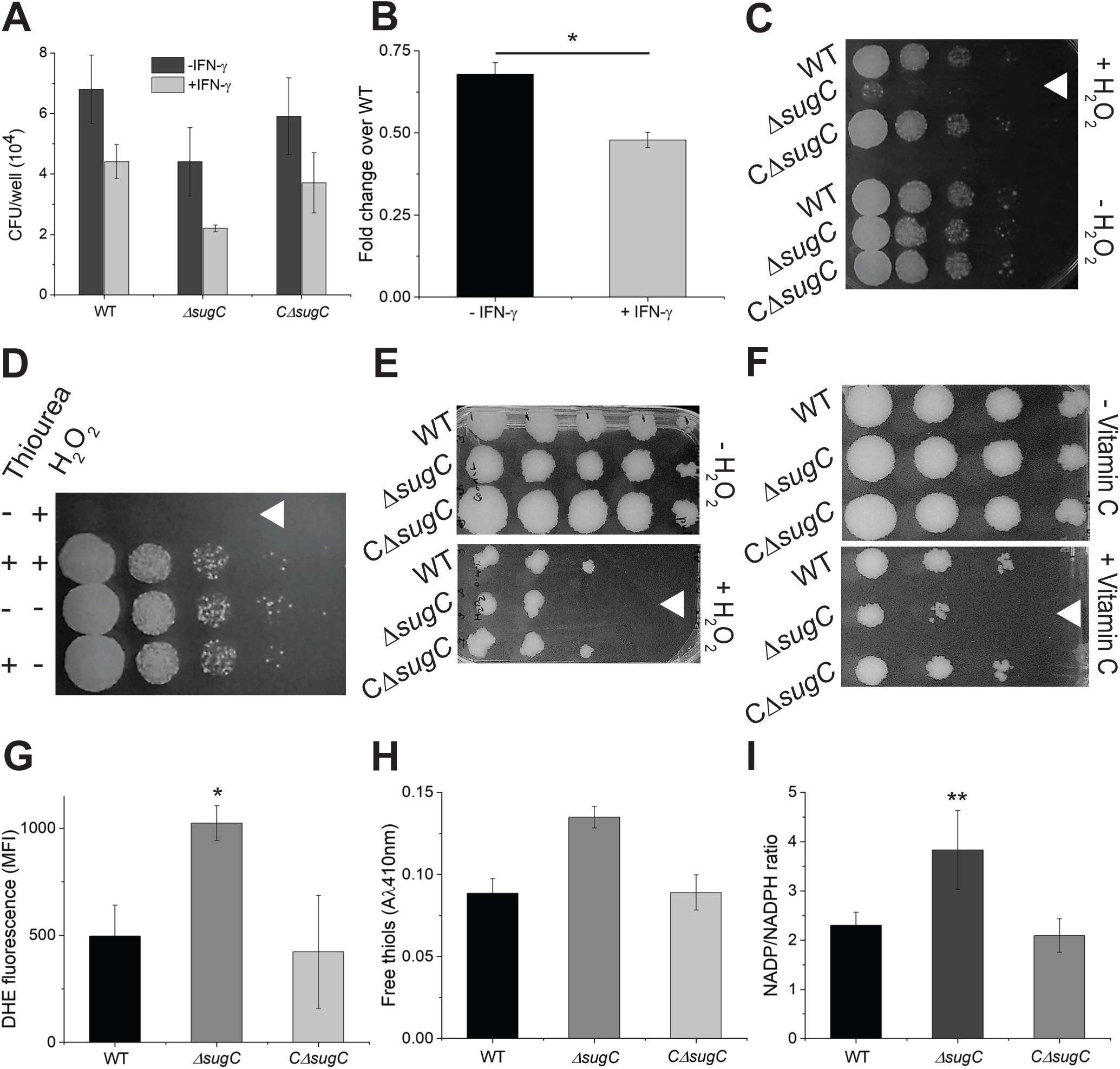
Loss of trehalose recycling upsets mycobacterial redox balance and *M. tuberculosis* survival in macrophages. (A) Survival of wild-type, *ΔsugC* and complemented *ΔsugC* (C*ΔsugC*), *M. tuberculosis* (*Mtb*) in immortalized C57BL/6 bone marrow-derived macrophages +/- IFN-γ treatment at 3 days post-infection. Multiplicity of infection, 1:5 (macrophages:*M. tuberculosis*). Experiment was performed at least three times either in duplicate or triplicate; one representative experiment shown. (B) Survival of *ΔsugC* relative to wild-type *M. tuberculosis* +/- IFN-γ. Ratios plotted from three independent experiments, including data from Figure 3A. (C) and (D), Sensitivity of carbon-deprived *M. smegmatis* or (E) and (F), *M. tuberculosis* to H_2_O_2_ or vitamin C. 10-fold serial dilutions were plated at the indicated time points. White triangles highlight most sensitive strain or condition. Experiments were performed at least three times; representative data are shown. (D), Effect of thiourea pre-treatment on H_2_O_2_ sensitivity of *ΔsugC M. smegmatis*. (G) Staining of *M. smegmatis* cultured in 0.02% glucose-supplemented medium by superoxide indicator dye dihydroethidium (DHE). Fluorescence detected by flow cytometry and median fluorescence intensity (MFI) plotted. Experiment was performed three times in triplicate; one representative experiment shown. (H) Quantification of total free thiols. Cell lysates of *M. smegmatis* that had been cultured in 0.02% glucose-supplemented medium were incubated with 5,5′-dithiobis (2-nitrobenzoic acid) and the absorbance at 412 nm was measured. Experiment was performed two times in triplicate with similar results; one experiment shown. (I) NADP^+^/ NADPH levels from *M. smegmatis* cultured in 0.02% glucose-supplemented medium. Ratios plotted from three independent experiments performed in triplicate. Error bars, standard deviation. Statistical significance compared to fold change in CFU of *ΔsugC* relative to wild-type in - IFN-γ and + IFN-γ, (B), and DHE fluorescence and NADP/NADPH ratio of *ΔsugC* and wild-type grown in 0.02% glucose, (G) and (I), was assessed by two-tailed Student’s t test. *, p<0.05; **, p<0.005.

### Loss of trehalose recycling alters mycobacterial redox balance

Overexpression of the LpqY component of the LpqY-SugABC transporter tolerizes *M. smegmatis* to a variety of antibiotics, and conversely, loss of the gene sensitizes *M. tuberculosis* to these drugs (Danelishvili et al., 2017). Susceptibility to many antibiotics has been linked to ROS overproduction, although the nature of this relationship is still unclear (Liu and Imlay, 2013; Van Acker and Coenye, 2017). One of the many consequences of IFN-γ activation is to enhance the production of (myco)bactericidal reactive oxygen species (ROS) in the macrophage (Hu et al., 2019). Given that the *ΔsugC* growth defect moderately worsened when macrophages were activated with IFN-γ (**Figures 3A, 3B**), we asked whether blocking trehalose recycling disrupts redox homeostasis. We focused on testing this hypothesis under growth-limiting (**Figure S1**, (Yang et al., 2014)) carbon limitation as trehalose recycling is enhanced in this condition (**Figures 2D, 2E**).

*ΔsugC M. smegmatis* and *M. tuberculosis* were more susceptible to H_2_O_2_ than wild-type (**Figures 3C, 3E**). Reintroduction of *sugC* (**Figure 3C, 3E**) or the addition of the ROS scavenger thiourea reversed this sensitivity (**Figure 3D**). *ΔsugC M. tuberculosis* was also more susceptible to vitamin C (**Figure 3F**), the mycobactericidal effect of which depends on ROS production (Vilcheze et al., 2013). Loss of trehalose recycling enhanced the fluorescence of dihydroethidium (DHE), a superoxide indicator dye (**Figure 3G**, (Owusu-Ansah et al., 2008)). Propidium iodide staining remained unchanged (**Figure S4E**), however, suggesting that the effect was not due to nonspecific differences in uptake, efflux or cell size.

Mycothiol (MSH) is the primary, cysteine-containing antioxidant in Actinobacteria (Newton et al., 2008; Ung and Av-Gay, 2006) and is the major source of free thiols in the mycobacterial cytoplasm (Kumar et al., 2011). The total free thiol content was modestly enhanced in the absence of *sugC* and restored to wild-type levels upon complementation (**Figure 3H**). We hypothesized that the increase in free, cytoplasmic thiols in the *sugC* mutant might be an adaptation to counteract the higher basal levels of superoxide. MSH is oxidized to mycothiol disulfide (Newton et al., 2008) and regeneration of MSH by mycothiol disulfide reductase (Mtr) requires NADPH (Kumar et al., 2011). Consistent with a drive to maintain a reduced mycothiol pool, we observed increased NADP:NADPH (**Figure 3I**) in *ΔsugC M. smegmatis.* Taken together, our data suggest that trehalose recycling that occurs during carbon limitation (**Figures 2D, 2E**) is necessary for redox homeostasis.

### Depletion of trehalose pool or inhibition of trehalose catabolism is not sufficient to cause oxidative stress

Trehalose in plants, fungi and bacteria can protect against ROS (Kuczynska-Wisnik et al., 2015; Lunn et al., 2014; Luo et al., 2008; Wang et al., 2018). We wondered whether the ROS phenotypes of *ΔsugC* mycobacteria in carbon-limited medium (**Figure 3**) were the direct result of a diminished cellular pool of trehalose. In addition to the LpqY-SugABC salvage pathway, there are several catabolic and anabolic pathways for trehalose (**Figure 4A**): OtsA and OtsB convert phosphorylated glucose intermediates to trehalose; TreY and TreZ degrade the glucose polymer α**-**glucan into trehalose; TreS converts trehalose to maltose; trehalase degrades trehalose into glucose. OtsAB is the primary source of trehalose in wild-type mycobacteria (Murphy et al., 2005). The free, cytoplasmic trehalose pool decreased ∼2-3-fold in the absence of *sugC* or *otsA* (**Figure 4B**) or overexpression of trehalase (**Figure S5D**), but was unchanged in *ΔtreYZ* (**Figure 4B**). The *ΔsugC* and *tre* overexpression strains were more sensitive to H_2_O_2_ than wild-type (**Figures 3C, 3E, S5C**). However, *ΔotsA M. smegmatis* did not have ROS phenotypes (**Figures 4C, 4D**) despite having the lowest amount of intracellular trehalose (**Figure 4B**). Thus, trehalose depletion alone is not sufficient to cause oxidative stress.

**Figure 4.**
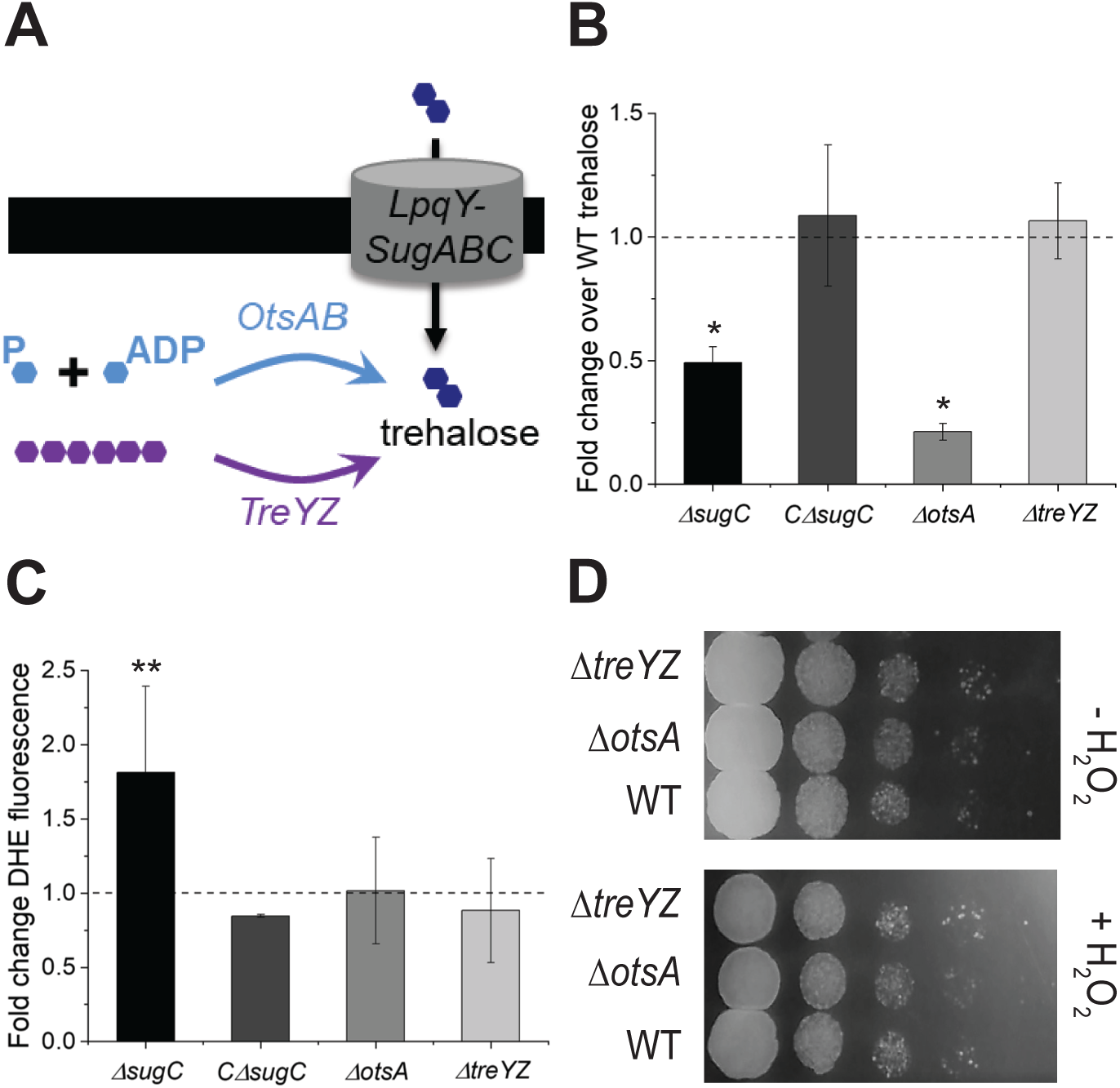
Depletion of trehalose pool or inhibition of trehalose catabolism is not sufficient to cause oxidative stress. (A) Cartoon showing *de novo* trehalose synthetic pathways. Light blue, phosphorylated glucose intermediates. Purple, α-glucan polymer. (B) Quantification of intracellular trehalose from *M. smegmatis* cultured in 0.02%-supplemented medium. Data plotted from three independent experiments performed in triplicate. The obtained values for colorimetric detection of trehalose were normalized to wild-type. (C) Staining of *M. smegmatis* cultured in 0.02% glucose-supplemented medium by superoxide indicator dye dihydroethidium (DHE). Fluorescence detected by flow cytometry and median fluorescence intensities (MFI) of the different mutants were normalized to wild-type. Data plotted from two independent experiments for complemented *ΔsugC (*C*ΔsugC*), and from four independent experiments for the rest of the strains. Experiments performed in triplicate. DHE fluorescence for all strains normalized to that of wild-type strain. (D) Sensitivity of carbon-deprived wild-type, *ΔotsA* and *ΔtreYZ M. smegmatis* to H_2_O_2_. 10-fold serial dilutions were plated at the indicated time points. Experiment was performed three times; representative data shown. Error bars, standard deviation. Statistical significance compared to wild-type was assessed by two-tailed Student’s t test. *, p<0.05; **, p<0.005.

In mature *M. tuberculosis* biofilms, trehalose is shunted into glycolytic and pentose phosphate intermediates in a TreS-dependent manner (Lee et al., 2019). Loss of TreS results in depletion of NADPH and of the NADPH-requiring antioxidant precursor γ-glutamylcysteine, which in turn heightens sensitivity to ROS. These data suggest that catabolism of recycled trehalose by TreS or trehalase might mitigate ROS. Consistent with their catalytic activities, loss of the enzymes resulted in slightly elevated levels of trehalose (**Figure S5A**). However, neither *ΔtreS* nor *Δtre* was as sensitive to H_2_O_2_ as *ΔsugC* (**Figure S5B**) under our conditions. These data suggest that enhanced ROS in the absence of trehalose recycling is either not a consequence of blocking downstream catabolism, or that it requires loss of both enzymes. Given our observation that trehalase overexpression potentiates ROS sensitivity (**Figure S5C**), we favor the former model.

### OtsA is required for oxidative stress when trehalose pool is depleted

A possible endogenous source of superoxide in the bacterial cell is ATP-generating oxidative phosphorylation. One method for estimating flux through this pathway is by O_2_ consumption. We observed more methylene blue decolorization for *ΔsugC* in carbon-limited medium (**Figure 5A**), indicating that respiration is enhanced in the absence of trehalose recycling. Notably, however, the mutant had lower levels of ATP than wild-type (**Figure 5B**). These data are consistent with a model in which ATP depletion in the absence of trehalose recycling drives oxidative phosphorylation, and in turn, ROS production. Alternatively, or additionally, redox stress might be a secondary consequence of metabolic perturbation.

**Figure 5.**
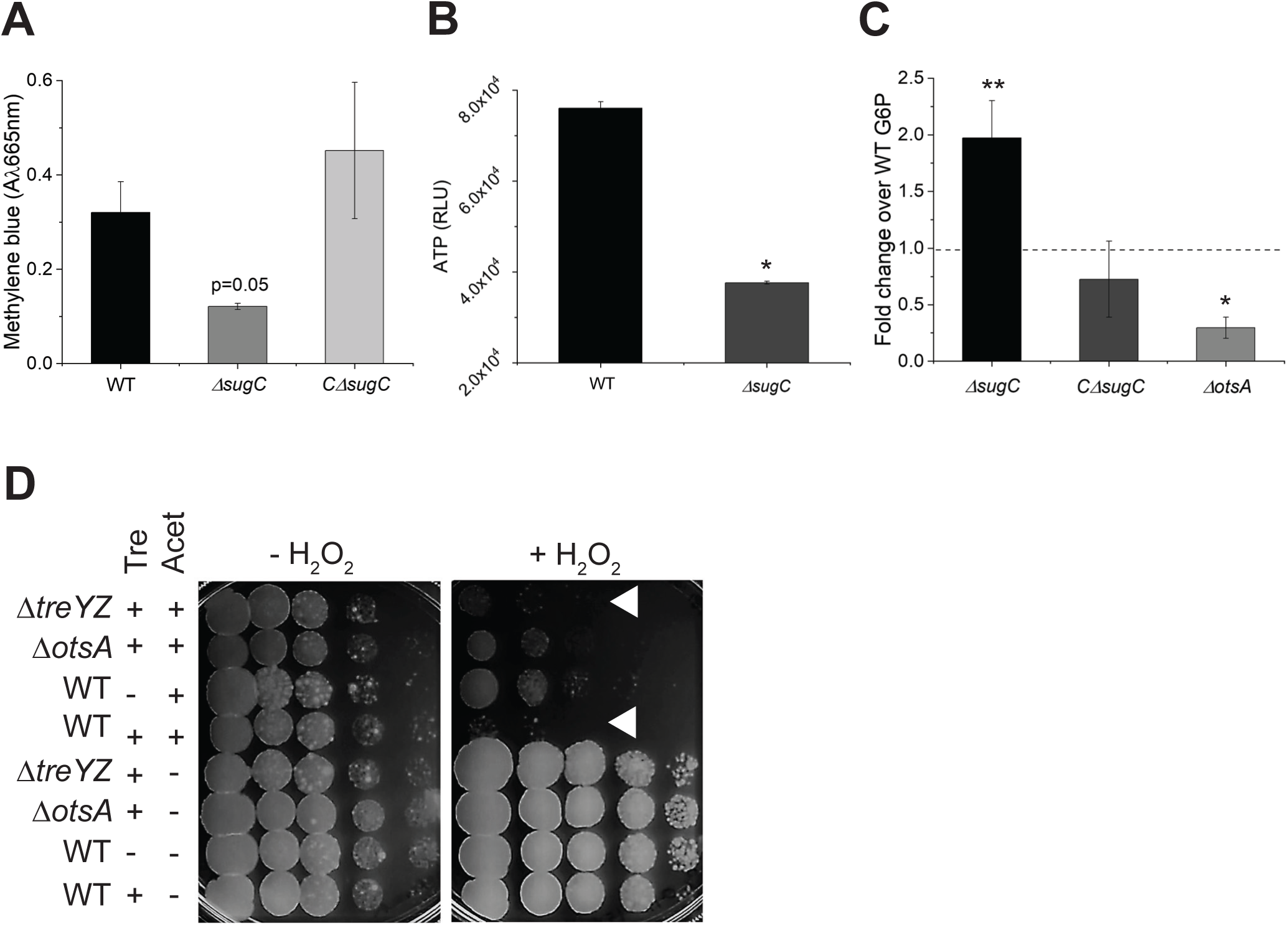
OtsA is required for oxidative stress when trehalose pool is depleted. (A) Oxygen consumption of *M. smegmatis* cultured in 0.02% glucose-supplemented medium. Strains were incubated +/- methylene blue and absorbance at 665 nm was measured. Absorbance from untreated samples subtracted. Experiment was performed three times in triplicate; one representative experiment shown. (B) ATP levels of *M. smegmatis* cultured in 0.02% glucose-supplemented medium. Protein concentration-normalized cell lysates were incubated with BacTiter-Glo™ reagent and luminescence was measured in relative light-forming unites (RLU). Experiment was performed at least three times in triplicate; one representative experiment shown. (C) Glucose-6-phosphate (G6P) levels of *M. smegmatis* cultured in 0.02% glucose-supplemented medium. Protein concentration-normalized cell lysates were incubated with G6P working solution and G6P level was measured in 96 well plate by monitoring the absorbance ratio at A575nm/A605nm. Data are plotted for three independent experiments performed in duplicate. G6P levels normalized to those of wild-type. (D) Sensitivity of carbon-deprived *M. smegmatis* to H_2_O_2_. 10-fold serial dilutions were plated at the indicated time points. White triangles highlight most sensitive strains or conditions. –Tre, plasmid backbone only; +Tre, plasmid with gene encoding trehalase under acetamide-inducible promoter; Acet, acetamide. Experiment was performed at least three times; representative data shown. Error bars, standard deviation. Statistical significance compared to wild-type and *ΔsugC* strains grown under 0.02% glucose supplemented 7H9 medium (A, B and D), or wild-type and *ΔotsA* strains grown under 0.02% glucose supplymented 7H9 medium (D), was assessed by two-tailed Student’s t test. *, p<0.05; **, p<0.005.

Why might carbon-starved, slow-growing mycobacteria consume ATP in the absence of trehalose recycling? We noted that mycomembrane synthesis continues unabated in *ΔsugC* (**Figure S4C**) and that TMM remains at wild-type levels (**Figures S4A, S4B**). The synthetic lethal interactions between *otsA* and *treYZ* or *lpqY*-*sugAC* in *M. tuberculosis* (Korte et al., 2016) suggest functional redundancy between the pathways encoded by these genes. The TreYZ pathway does not require energy to break down α**-** glucan into trehalose but OtsA and OtsB convert phosphorylated glucose intermediates to trehalose. In glucose-limited conditions, moreover, trehalose synthesis via the OtsAB pathway likely requires additional ATP to drive gluconeogenesis. We wondered whether oxidative stress that occurs in the absence of LpqY-SugABC or upon trehalase overexpression (**Figures 3, S5B**) might be from induction of the OtsAB pathway. The end product of gluconeogenesis is glucose-6-phosphate (G6P). The levels of G6P were elevated in *ΔsugC* but suppressed in *ΔotsA* (**Figure 5C**), respectively consistent with higher and lower levels of gluconeogenesis. We next wished to test whether induction of the OtsAB pathway upsets redox balance in carbon-deprived mycobacteria. Given the synthetic lethal interaction between *sugC* and *otsA* (Korte et al., 2016), we opted to deplete the trehalose pool by inducible trehalase overexpression. We compared the H_2_O_2_ sensitivity of strains that overexpress trehalase in wild-type, *ΔotsA* and *ΔtreYZ* backgrounds. Loss of OtsA, but not TreYZ, partially rescued sensitivity of *M. smegmatis* to H_2_O_2_ upon trehalase overexpression (**Figure 5D**). Taken together, our results support the idea that maintenance of the trehalose pool via the energy-intensive OtsAB pathway leaves carbon-starved, slow-growing mycobacteria vulnerable to oxidative stress.

## Discussion

Hints of mycomembrane plasticity began to appear in the early 1900s, when it was recognized that acid-fastness–a hallmark staining property still used for microscopy-based diagnosis of *M. tuberculosis*—varied with nutrient supply (Cadena et al., 2017; Seiler et al., 2003; Vilcheze and Kremer, 2017). More recent work supports the idea that the mycomembrane is reconfigured *in vivo* and in response to host-mimicking stresses (Bacon et al., 2014; Baker and Abramovitch, 2018; Bhamidi et al., 2011; Eoh et al., 2017; Galagan et al., 2013; Kieser and Rubin, 2014; Ojha et al., 2010; Shui et al., 2007; Yang et al., 2014). The mechanisms by which these cell surface alterations occur are still emerging but have been attributed primarily to catabolic pathways (Eoh et al., 2017; Yang et al., 2014). We took advantage of recent advances in metabolic labeling (Foley et al., 2016; Siegrist et al., 2015) to show that mycomembrane remodeling under carbon deprivation also involves anabolic reactions (**Figure 1C**), a counterintuitive result as mycobacterial replication (**Figures 1C, S1B, S1C**) and presumably overall metabolic activity are sluggish. Our data collectively indicate that the net result of such reactions is decreased TDM and spatial rearrangement of AGM. We previously showed that synthesis of peptidoglycan along the non-expanding sidewall of *M. smegmatis* is enhanced in response to cell wall damage (Garcia-Heredia et al., 2018). AGM synthesis under carbon starvation also occurs along the cell periphery (**Figure 1F**), further supporting the notion that mycobacteria can edit their cell surface in a growth-independent fashion.

The adaptive consequences of mycomembrane remodeling are manifold (Karakousis et al., 2004; Nobre et al., 2014; Rajni et al., 2011). For example, bulk decreases in TDM and AGM abundance are known to increase mycobacterial cell permeability, which in turn enhances nutrient uptake and antimicrobial susceptibility (Gebhardt et al., 2007; Jackson et al., 1999; Yang et al., 2014). Although we do not observe gross changes in the amount of AGM under nutrient deprivation (**Figure 1D**), the primary site of synthesis shifts from the pole to sidewall (**Figure 1F**). The concomitant reduction in permeability (**Figure 1E**)—despite an overall decrease in TDM abundance—suggests that the subcellular distribution of AGM also contributes to the barrier function of the mycobacterial cell envelope. Beyond enabling edits to the structural components of the mycomembrane, remodeling reactions liberate smaller molecules that influence cell physiology. Free trehalose released by TDM and AGM synthesis can be recycled into glycolysis or pentose phosphate intermediates, or act as a stress protectant or compatible solute in the cytoplasm (Eoh et al., 2017; Lee et al., 2019; Nobre et al., 2014; Shleeva et al., 2017). Our data suggest that it can also be directly refashioned into trehalose-containing, cell surface glycolipids (**Figures 2D, 2E**), likely TMM. Free mycolic acids generated by TDM hydrolysis are components of biofilm matrix (Ojha et al., 2010) and, like trehalose, serve as carbon sources (Rafidinarivo et al., 2009). We speculate that they may additionally be reused together with recycled trehalose to make TMM.

How do mycobacteria power mycomembrane remodeling when faced with a loss of nutrients? The three isoforms of the TMM-consuming Antigen 85 complex, encoded in *M. tuberculosis* by *fbpA, fbpB* and *fbpC*, have partially redundant acceptor specificities (Jackson et al., 1999; Puech et al., 2002). However only *fbpC* is upregulated in nutrient-starved *M. tuberculosis* (Betts et al., 2002; Jamet et al., 2015) making Ag85C an obvious candidate for performing synthetic reactions under that condition. Perhaps the more interesting question, however, is the source of the energetically-expensive TMM building blocks. Breakdown of TDM by TDMH furnishes free mycolic acids and TMM, the latter of which could serve as a donor for sidewall AGM synthesis (Holmes et al., 2019; Ojha et al., 2010). While such a pathway would not require ATP, it would be limited by the amount of TDM loss that can be tolerated without lysis (Ojha et al., 2010; Yang et al., 2013) or reduced resilience to host stress (Yang et al., 2014). Our data suggest that *M. smegmatis* and *M. tuberculosis* also generate TMM in the cytoplasm from recycled trehalose (**Figures 2D, 2E**). An intracellular route of TMM generation would limit TDM loss, thereby preserving mycomembrane integrity. Use of recycled materials in turn would allow the mycobacterial cell to reap the benefit(s) of sidewall AGM fortification while minimizing energy expenditure. In the absence of trehalose recycling, *de novo* synthesis supplies the sugar and mycomembrane remodeling continues unabated (**Figure S4**). The cost of from-scratch, OtsAB-mediated anabolism is not apparent under standard *in vitro* culture conditions but sensitizes *M. smegmatis* and *M. tuberculosis* to ROS (**Figure 3**) and may contribute to defective *M. tuberculosis* growth during infection (**Figure 3A,** (Kalscheuer et al., 2010b)).

Trehalose is a cytoplasmic stress protectant and compatible solute and, in many types of bacteria, a carbon source (Elbein et al., 2003; Kuczynska-Wisnik et al., 2015; Zhao et al., 2019). Mycobacteria and related organisms are relatively unique in using trehalose for extracellular purposes, to build their outer cell envelope. As the sugar fluxes in and out of central metabolism and the mycomembrane via several synthetic (OtsAB, TreYZ) and degradative (TreS, trehalase) processes, trehalose utilization may be particularly vulnerable to perturbations that induce redox and metabolic imbalances. Like carbon-limited *ΔsugC M. smegmatis* or *M. tuberculosis*, biofilm cultures of *ΔtreS M. tuberculosis* are more susceptible to ROS. However, our data suggest that the mechanisms are distinct. In mature biofilms, trehalose is shunted away from TMM and TDM synthesis into glycolytic and pentose phosphate intermediates in a TreS-dependent manner (Lee et al., 2019). By contrast, we find that TMM levels are maintained during the time frame of our experiment, either by LpqY-SugABC, in wild-type organisms, or by *de novo* synthesis, in *ΔsugC* mutants. While biofilm *ΔtreS M. tuberculosis* are likely more sensitive to ROS because they are depleted for the antioxidant precursor γ-glutamylcysteine (Lee et al., 2019), carbon-limited *ΔsugC M. smegmatis* have higher levels of ROS-counteracting, cytoplasmic thiols (**Figures 3G, 3H**). These and other metabolite data are most consistent with the idea that enhanced ROS production and susceptibility (**Figure 3**) in the absence of trehalose recycling stems from increased anabolism of the sugar rather than decreased catabolism. While we focus here on mycomembrane remodeling that occurs within 1-3 days of adaptation to carbon-limited medium, the TreS-dependent, trehalose-catalytic shift occurs in 4-5-week-old biofilms. Under our conditions, loss of TreS has no impact on ROS susceptibility (**Figure S5B**). While we cannot rule out stress- or species-specific differences between the two studies, we favor a model in which the adaptive role of trehalose changes over time: early fortification of the cell envelope, to protect against immediate environmental insults, and later rewiring of central carbon metabolism, to maintain ATP and antioxidant levels.

The presence of a retrograde transporter enables trehalose to cycle in and out of the cell and serve as a metabolic node between the mycomembrane and cytoplasm. Recycling of the sugar is well-known to enhance *M. tuberculosis* survival in a mouse model of tuberculosis. It is widely hypothesized that the *in vivo* growth defects of trehalose recycling mutants stem from progressive carbon starvation (Kalscheuer and Koliwer-Brandl, 2014; Kalscheuer et al., 2010b; Nobre et al., 2014). At later time points, nutrient deprivation coupled with loss of trehalose catabolism may indeed reduce fitness. However, our data suggest a more complex model for early adaptation to stress, namely that *de novo* synthesis induced in the absence of recycling consumes ATP and increases respiration and ROS sensitivity (**Figure 6**). The metabolic and redox phenotypes of a trehalose recycling mutant resemble those elicited by futile cycling (Adolfsen and Brynildsen, 2015; Brynildsen et al., 2013) or certain bactericidal antibiotics (Yang et al., 2017), and indeed, loss of trehalose recycling sensitizes *M. tuberculsosis* to multiple drugs (Danelishvili et al., 2017). There is growing appreciation that processes downstream of target inhibition, particularly metabolic and redox dysfunction, contribute to the cidality of certain antibiotics (Yang et al., 2017). We posit that analogous dysfunction triggered by forced *de novo* synthesis of energy-expensive macromolecules could be a fruitful avenue for potentiating immune and antibiotic activity against bacterial pathogens, including those that inhabit growth-limiting, nutrient-deprived host niches.

**Figure 6.**
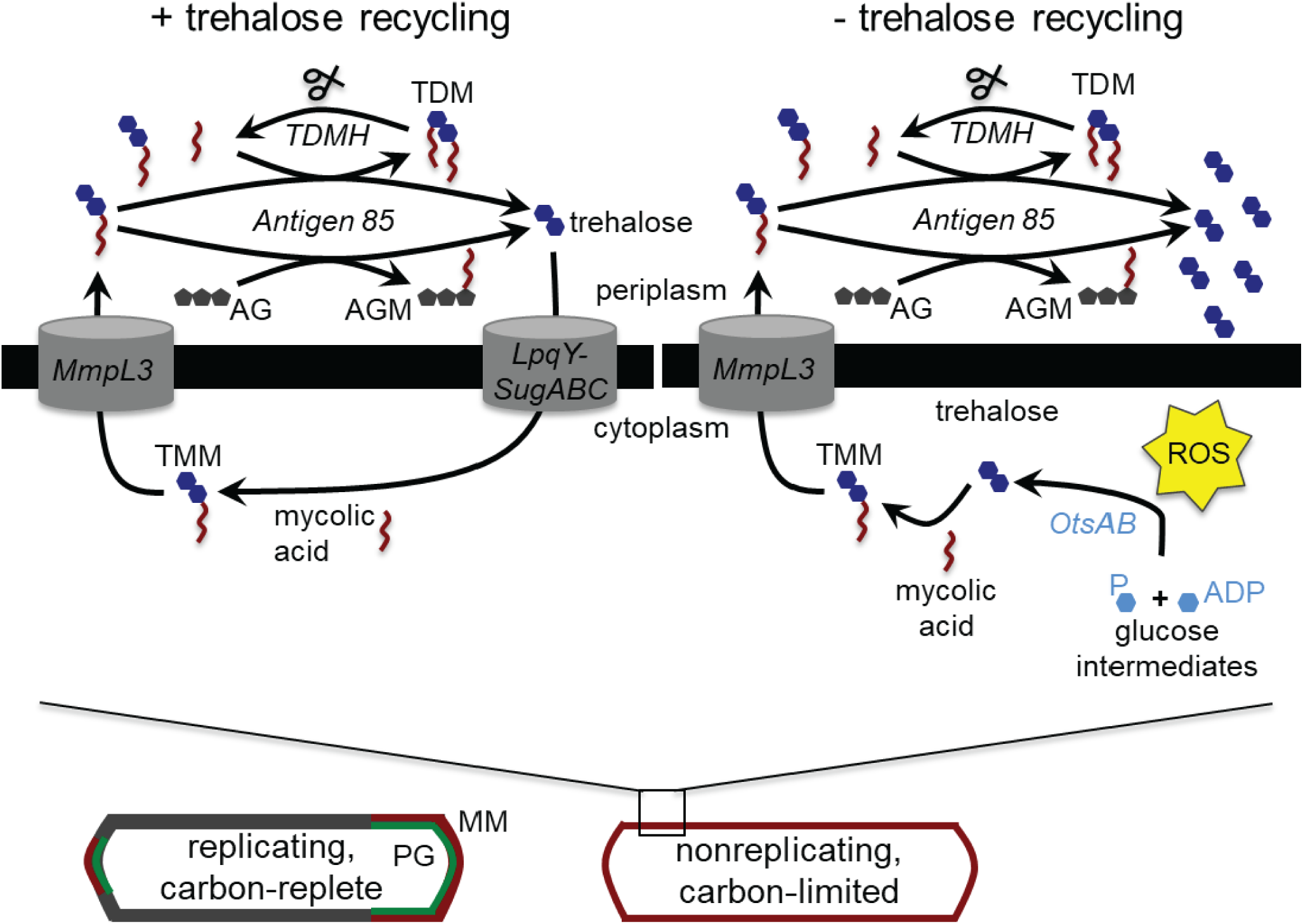
Model for the role of trehalose recycling in mycomembrane remodeling under nutrient limitation. Bottom, mycobacteria respond to growth-limiting carbon deprivation by turning over TDM and synthesizing AGM along the entire cell periphery. Top left, in wild-type cells, the TMM building blocks are obtained at least in part from trehalose recycled by LpqY-SugABC. Top right, in mutants unable to recycle trehalose, TMM is supplied by *de novo* trehalose synthesis. Increased flux through the OtsAB pathway depletes ATP, drives respiration and confers ROS sensitivity.

## Acknowledgments

We are grateful to Drs. Rainier Kalscheuer for wild-type, *ΔsugC* and complemented *ΔsugC M. tuberculosis,* Christopher Sassetti for immortalized bone marrow-derived macrophages, Yasu Morita for assistance with thin layer chromatography and Hyungjin Eoh for thoughtful feedback on the manuscript. Research was supported by NIH DP2 AI138238 (M.S.S.) and NSF CAREER 1654408 (B.M.S.).

## Author Contributions

A.A.P. Conceptualization, methodology, validation, formal analysis, investigation, visualization, project administration, data curation, writing – reviewing and editing. C.R.C. Validation, investigation, writing – reviewing and editing, data curation, software. J.G. Supervision, writing – reviewing and editing. B.M.S. writing – Methodology, writing – reviewing and editing, resources, funding acquisition. M.S.S. Conceptualization, methodology, visualization, resources, writing – Original Draft, reviewing and editing, supervision, project administration, funding acquisition.

## Supplementary Figure Legends

**Figure S1. Effect of carbon source on cell envelope labeling.**

(A) O-AlkTMM and (B) HADA labeling of *M. smegmatis* cultured in medium supplemented with different carbon sources. Alkynes were detected by CuAAC reaction with carboxyrhodamine 110 azide. Data are normalized to labeling in 2% glucose-supplemented medium and plotted from three independent experiments. Cyan autofluorescence subtracted from values in (B).

(C) Growth profile of *M. smegmatis* grown in different carbon sources. Experiment was performed twice in triplicate with similar results; one experiment shown.

Error bars, standard deviation.

Related to **Figure 1**.

**Figure S2. Quantitation of mycomembrane components.**

Representative images of thin-layer chromatography (TLC) of (A) trehalose dimycolate (TDM), (B) free mycolic acids from culture supernatant (CS-MA) or (C) covalently-bound mycolic acids (AGM-MA). Wild-type (WT) and *ΔsugC* strain were cultured for 24 hours in 0.02% or 2% glucose-supplemented medium and processed for lipid extraction. Samples were normalized by wet pellet weight ((A), (C)) or by optical density (B). STD, standard. For (A), chloroform:methanol:acetone (90:15:10) solvent system was used. TLC was developed by spraying with 5% H_2_SO_4_ in ethanol and then charring the plate at 110°C for 10 minutes. For (B), chloroform:methanol (96:4) solvent system was used. TLC was developed by spraying 5% molybdophosphoric acid in ethanol and charring the plate at 110°C for 10 minutes. For (C), the extracted mycolic acids-arabinogalactan-peptidoglycan (mAGP) was used for MA extraction by alkaline hydrolysis.

Related to **Figure 1**.

**Figure S3. Metabolic labeling and quantitation of TMM.**

(A) Metabolic labeling of wild-type and *ΔsugC M. smegmatis* in 0.02% and 2% glucose-supplemented medium with 6-TreAz (labels TMM). After fixation, alkynes were detected by CuAAC reaction with carboxyrhodamine 110 alkyne.

(B) & (C) Representative images of thin-layer chromatography (TLC) of TMM from wild-type (WT), *ΔsugC* and complemented (*CΔsugC*) *M. smegmatis* (B) or *M. tuberculosis* (C) cultured for 24 hours in low or high carbon medium. Samples were normalized by wet pellet weight. TLC was run in the chloroform:methanol:H_2_O (80:20:2) solvent system and developed by spraying with 5% H_2_SO_4_ in ethanol and then charring the plate at 110°C for 10 minutes. STD, standard.

Related to **Figure 2**.

**Figure S4. Mycomembrane reorganization under carbon deprivation can occur in the absence of trehalose cycling.**

(A) and (B), quantitation of TMM abundance in *ΔsugC* and complemented (*CΔsugC*) *M. smegmatis* (B) or *M. tuberculosis* (C) cultured for 24 hours in low or high carbon medium. TLCs were scanned and processed in ImageJ (Schneider et al., 2012). Data were normalized to wild-type and plotted from three independent experiments, including images from **Figure S3B** and **S3C**.

(C) Metabolic labeling of wild-type and *ΔsugC M. smegmatis* in 0.02% and 2% glucose-supplemented medium with O-AlkTMM (primarily labels AGM), N-AlkTMM (labels TDM) and alkDala (labels peptidoglycan). After fixation, alkynes were detected by CuAAC reaction with carboxyrhodamine 110 azide. Fluorescence was quantitated by flow cytometry and expressed as median fluorescence intensity (MFI).

(D) TDM turnover under nutrient deprivation. *ΔsugC M. smegmatis* cultured in 0.02% glucose-supplemented medium was labeled with metabolic probes O-AlkTMM, N-AlkTMM or HADA (labels peptidoglycan), then washed. Aliquots were taken at the indicated time points. After fixation, alkynes were detected by CuAAC reaction with carboxyrhodamine 110 azide. Fluorescence was quantitated by flow cytometry, with the median fluorescence intensities normalized to the initial, 0 h time point for each probe.

(E) Propidium iodide staining of wildtype and *ΔsugC M. smegmatis* upon adaptation to 0.02% glucose-supplemented medium. Fluorescence was quantitated by flow cytometry and median fluorescence intensity (MFI) was plotted. Experiment was performed three times in triplicate; one representative experiment shown.

**Figure S5. Intracellular trehalose abundance and ROS sensitivity upon modulation of trehalose catabolism.**

(A) Quantification of intracellular trehalose from wild-type, *Δtre* and *ΔtreS M. smegmatis* cultured in 0.02% glucose-supplemented medium. Experiment was performed three times in triplicate; one representative experiment is shown.

(B) and (C), H_2_O_2_ sensitivity of wild-type, *ΔsugC, Δtre* and *ΔtreS* (B) or *tre*-overexpressing (C) *M. smegmatis* cultured in 0.02% glucose-supplemented medium. 10-fold serial dilutions were plated at the indicated time points. White triangles highlight most sensitive strains or conditions. –Tre, plasmid backbone pYAB-EV only; +Tre, plasmid with gene encoding trehalase (pYAB-Tre) under acetamide-inducible promoter; Acet, acetamide. Experiments were performed 2-3 times; representative data shown.

(D) *tre* overexpression drains intracellular pool of trehalose. Ratios of trehalose from acetamide-induced:uninduced for *M. smegmatis* expressing pYAB-EV and pYAB-Tre. Data plotted from three independent experiments performed in triplicate.

Error bars, standard deviation. Statistical significance was assessed by two-tailed Student’s t test. *, p<0.05.

Related to **Figure 4**.

