## Supplementary data for "Trehalose recycling promotes energy-efficient mycomembrane reorganization in nutrient-limited mycobacteria"

Running title: Trehalose recycling supports redox homeostasis

#### Materials and Methods

**Bacterial strains and culture conditions.** *M. smegmatis* mc$^2$155 was grown in Middlebrook 7H9 growth medium (HiMedia, India) supplemented with Tween-80 (7H9T) and glucose (2% or 0.02%) at 37°C unless otherwise specified in the text. Two day-old primary cultures of *M. smegmatis* grown in 2% glucose were normalized to an $OD_{600}$ of 0.1 in fresh 7H9T supplemented with 2% or 0.02% glucose and allowed to grow for 24 hours. *M. tuberculosis* H37Rv strains (gifts of Dr. Rainier Kalscheuer) were grown in Middlebrook 7H9 medium (BD Difco, Franklin Lakes, NJ) supplemented with Tween-80 and OADC (BD BBL, Sparks, MD). For starvation of *M. tuberculosis*, cultures grown in 7H9T-OADC to $OD_{600}$ 0.8-1.0 were collected by centrifugation and washed once with 7H9T (no OADC) and resuspended in 7H9T (starvation medium) to a normalized $OD_{600}$ of 1. To prepare a strain that expresses *tre*, the gene that encodes trehalase, under an acetamide-inducible promoter, we PCR amplified *tre* from genomic DNA of *M. smegmatis* by using 4535For_Acet (tgatgtgctctagagttctgcaacagaccgagcc) and 4535Rev_Acet (ggcctgatctagacatcggggcgttcgcgg) primers. The resulting PCR product was ligated in pYAB033 vector (gift of Dr. Yasu Morita) at XbaI site and transformed in *E. coli* XL-1 blue strain. The colonies were screened by colony PCR and the obtained plasmid was sequence-confirmed. Bacteria used in this study are listed in **Table S1.**

**ROS sensitivity.** *M. smegmatis* grown in 0.02% glucose for 24 hours were normalized to $OD_{600}$ of 1. The cultures were then treated with 0.15% $H_2O_2$ for 10 minutes at 37°C with shaking. The trehalase overexpression strains were grown for 20 hours in 0.02% glucose and then induced with 0.2% acetamide for an additional 10 hours before being treated with 0.1% $H_2O_2$ for 10 minutes at 37°C with shaking. After $H_2O_2$ treatment, 3 μL of 10-fold serial dilutions made in PBS was spotted on 7H9-2% glucose agar. For thiourea rescue experiment, cultures were pretreated with 50 mM thiourea for 45 minutes prior to $H_2O_2$. For *M. tuberculosis*, cultures in starvation medium were grown for 5 days, normalized to $OD_{600}$ 0.1 in fresh starvation medium then treated with 0.4% of $H_2O_2$ for 2 hours at 37°C with shaking. After $H_2O_2$ treatment, 5 μL of 10-fold serial dilutions made in PBS were spotted on 7H10-OADC agar plate. For the vitamin C experiment, *M. tuberculosis* cultures in starvation medium were normalized to $OD_{600}$ 0.1 in fresh starvation medium. The cultures were then treated with 20 mM vitamin C for 2 days. After vitamin C treatment, 5 μL of 10-fold serial dilutions made in PBS were spotted on 7H10-OADC agar.

**Macrophage infections.** Immortalized C57BL/6 bone marrow-derived macrophages (iBMDM, gift of Dr. Christopher Sassetti) were seeded at $10^5$ cells/well in 24-well tissue culture plate and incubated at 37°C overnight. *M. tuberculosis* were added at 5:1 multiplicity of infection (MOI; bacteria:iBMDM) and incubated for 4 hours. After incubation the co-culture was washed twice with high glucose Dulbecco's modified Eagle's medium (DMEM, Genesee Scientific, San Diego, CA), to remove extracellular *M. tuberculosis,* and fresh DMEM-FBS-HEPES (5 mM) medium was added (FBS, Genesee Scientific, San Diego, CA and HEPES; Gibco, Paisley, PA, UK). IFN-γ (PeproTech, Rocky Hill, NJ) was added or not at 25 ng/mL concentration. The infected iBMDM were incubated for 3 days, then washed once with PBS and lysed with 0.05% Triton-X 100 in PBS. After lysis, 10 μL of 10-fold serial dilutions made in PBS were spotted on 7H10-OADC agar for determining colony-forming units (CFU).

**DHE staining.** *M. smegmatis* grown for 24 hours in 7H9T-0.02% glucose were normalized to $OD_{600}$ 1 with the same medium then treated with 5 μM dihydroethidium (DHE; Sigma, St. Louis, MO) for 30 minutes at 37°C. Fluorescence was analyzed by flow cytometry.

**Total thiol abundance.** The protocol for measuring the total thiol content was adopted from (Vilcheze et al., 2017). Briefly, 10 mL of *M. smegmatis* grown for 24 hours in 7H9T-0.02% glucose were centrifuged at 2500xg for 5 minutes, washed with buffer containing 50 mM Tris-Cl (pH 8) and 5 mM EDTA, and cell pellets were normalized by wet weight. Bacteria were resuspended in the same buffer and lysed by bead beating. Lysates were centrifuged at 16000xg for 15 minutes at 4°C and 5,5′-dithiobis (2-nitrobenzoic acid) was added to 100 μL of supernatants to a final concentration of 0.05 mM. Total thiol content was estimated by absorbance at λ412nm.

**Methylene blue.** *M. smegmatis* grown for 24 hours in 7H9T-0.02% glucose were adjusted to $OD_{600}$ 0.25. Cultures were split in two, one of which was treated with 0.005% methylene blue, then aliquoted to a 96-well plate. The plate was sealed with Microseal 'B' Adhesive Sealing Films (BioRad, UK) and incubated at 37°C for 4 hours with shaking. The seal was then removed and absorbance at λ665nm was measured. The difference between the λ665nm of treated and untreated samples was plotted.

**ATP, glucose-6-phosphate and NADP/NADPH quantitation**. ATP concentration was measured by by BacTiter-Glo (Promega, Madison, WI) luminescence kit. Glucose-6-phosphate (G6P) concentration and NADP/NADPH ratio were respectively measured with the Amplite™ (AAT Bioquest, Sunnyvale, CA) Colorimetric G6P Assay and Colorimetric NADP/NADPH Ratio Assay kits. *M. smegmatis* grown for 24 hours in 7H9T-0.02% glucose was washed once with PBS. The pellets were resuspended in PBS and lysed by bead beating. Lysates were normalized by total protein concentration using a BCA protein assay kit (Pierce, Rockford, IL) then processed according to the manufacturer's protocol.

**Trehalose quantitation**. For intracellular trehalose detection, *M. smegmatis* grown for 24 hours in 7H9T-0.02% glucose were washed once with PBS. Cell pellets were normalized by wet weight then resuspended in chloroform:methanol (1:1) for overnight incubation with shaking. The suspension was centrifuged at 10000xg for 5 minutes and the organic fraction was collected in a new tube. One part chloroform and one part water were added to the organic fraction and mixed vigorously in shaker for 15 minutes. Suspensions were centrifuged and the upper aqueous layers were processed per the manufacturer's instructions for the trehalose assay kit (Megazyme, Ireland). For extracellular trehalose detection, *M. smegmatis were* grown for 24 hours in 7H9T supplemented with 2% or 0.02% glycerol. Cultures were normalized to $OD_{600}$ 1 prior to centrifugation. The upper layer was collected and filtered through a 0.2 µM syringe. Filtrates were processed as above to detect trehalose.

**Lipid extraction and TLC.** For extractable lipid analysis, 10 mL of culture was washed with PBS and cell pellets were normalized by wet weight (*M. smegmatis*) or by $OD_{600}$ (*M. tuberculosis*). To obtain TDM and TMM, cell pellets were extracted with chloroform:methanol (2:1). The extracted lipids were separated by TLC (HPTLC silica gel, Millipore, Billerica, MA) with chloroform:methanol:acetone (90:15:10) and chloroform:methenol:$H_2O$ (80:20:2) for TDM and TMM, respectively (Foley et al., 2016; Touchette et al., 2017). 5% $H_2SO_4$ in ethanol was used to develop TLC.  Covalent mycolate extraction was adopted from (Payne et al., 2009). Briefly, mycolic-arabinogalactan-peptidoglycan (mAGP) complex was extracted from 100 mL of culture as described (Payne et al., 2009). The pellet was resuspended in PBS and sonicated to lyse the cells. Lysates were centrifuged and pellets were collected and washed with PBS. The pellets were resupended in 2% SDS in PBS and incubated at 80°C for 3 hours with intermediate shaking. They were then resuspended in 1% SDS, centrifuged, and washed twice with water, once with 80% acetone, and once with 100% acetone. Pellet were dried to obtain the final mAGP complex. Samples were normalized by mAGP weight, then resuspended in PBS + 0.05% Tween-80 (PBST) by water bath sonication. To extract mycolic acids from mAGP, the suspension was treated with 5% tetrabutylammonium hydroxide (TBAH) overnight with shaking. The extracted mycolic acids were seprated by treating with equal volume of dichloromethane followed by treatment with equal volume of 0.25 M HCl and water-washed as described (Payne et al., 2009). To extract free mycolic acids from culture supernatants, the $OD_{600}$ of *M. smegmatis* grown for 24 hours in 7H9T-2% or 0.02% glucose were normalized to 1 with 7H9T. The normalized cultures were centrifuged at 10000xg for 5 minutes and superntants were collected and passed through a 0.25 µm syringe filter. Supernatants (1 mL) were treated with 5% TBAH for 1 hour followed by an equal amount of dichloromethane and overnight incubation at room temperature with shaking. The suspension was then centrifuged at 10000xg and the lower organic layer was removed. The organic layer was evaporated and the pellet was mixed with 40 µL chloroform:methenol (2:1). Mycolic acids were separated by TLC using chloroform:methanol (96:4) as described (Ojha et al., 2010). 5% molybdophosphoric acid in ethanol was was used to develop the TLC.

**Propidium iodide (PI).** We assessed PI uptake as described (Sharma et al., 2015). Briefly, 50 µg/mL PI was added to *M. smegmatis* that had been cultured in 0.02% or 2% glucose. After incubating for 15 minutes at 37°C, samples were washed once with PBS and fluorescence was measured by flow cytometry.

**Cell envelope labeling.** Probes used in this study include alkDala (50 µM), HADA (500 µM), O-AlkTMM (50 µM), N-AlkTMM (250 µM) and 6-TreAz (50 µM). *M. smegmatis* labeling was performed mainly as described (Garcia-Heredia et al., 2018). Briefly, the $OD_{600}$ was normalized to 1 in the same medium. Cultures were shaken in the presence of probes for 30 min at 37°C for *M. smegmatis*. After incubation the cultures were washed twice with PBST and fixed or not with 2% formaldehyde at room temperature for 10 minutes. After fixation, cultures were washed with PBST. Alkynes were detected by CuAAC reaction with carboxyrhodamine 110 azide (Click Chemistry Tools, Scottsdale, AZ). Azides were detected on live, unfixed cells by SPAAC reaction with DBCO-Cy5 (Click Chemistry Tools, Scottsdale, AZ). Finally, cultures were washed thrice with PBST and fluorescence was measured by flow cytometery. For *M. tuberculosis,* the $OD_{600}$ for carbon-starved and unstarved cultures were normalized to 1 in the same media. Cultures were shaken in the presence of probes for 3 hours at 37°C then washed twice with PBST and subjected to SPAAC overnight at 37°C. Cultures were washed thrice with PBST and fixed with 4% formaldehyde overnight at room temperature prior to removal from the BSL3 facility.

**Microscopy analysis.** Fluorescence microscopy and image quantitation was performed exactly as described in (Garcia-Heredia et al., 2018).

#### Refrances

Foley, H.N., Stewart, J.A., Kavunja, H.W., Rundell, S.R., and Swarts, B.M. (2016). Bioorthogonal Chemical Reporters for Selective In Situ Probing of Mycomembrane Components in Mycobacteria. Angew Chem Int Ed Engl *55*, 2053-2057.

Garcia-Heredia, A., Pohane, A.A., Melzer, E.S., Carr, C.R., Fiolek, T.J., Rundell, S.R., Chuin Lim, H., Wagner, J.C., Morita, Y.S., Swarts, B.M.*, et al.* (2018). Peptidoglycan precursor synthesis along the sidewall of pole-growing mycobacteria. Elife *7*.

Ojha, A.K., Trivelli, X., Guerardel, Y., Kremer, L., and Hatfull, G.F. (2010). Enzymatic hydrolysis of trehalose dimycolate releases free mycolic acids during mycobacterial growth in biofilms. J Biol Chem *285*, 17380-17389.

Payne, K., Sun, Q., Sacchettini, J., and Hatfull, G.F. (2009). Mycobacteriophage Lysin B is a novel mycolylarabinogalactan esterase. Mol Microbiol *73*, 367-381.

Sharma, A., Pohane, A.A., Bansal, S., Bajaj, A., Jain, V., and Srivastava, A. (2015). Cell penetrating synthetic antimicrobial peptides (SAMPs) exhibiting potent and selective killing of *Mycobacterium* by targeting its DNA. Chemistry *21*, 3540-3545.

Touchette, M.H., Van Vlack, E.R., Bai, L., Kim, J., Cognetta, A.B., 3rd, Previti, M.L., Backus, K.M., Martin, D.W., Cravatt, B.F., and Seeliger, J.C. (2017). A Screen for Protein-Protein Interactions in Live Mycobacteria Reveals a Functional Link between the Virulence-Associated Lipid Transporter LprG and the Mycolyltransferase Antigen 85A. ACS Infect Dis *3*, 336-348.

Vilcheze, C., Hartman, T., Weinrick, B., Jain, P., Weisbrod, T.R., Leung, L.W., Freundlich, J.S., and Jacobs, W.R., Jr. (2017). Enhanced respiration prevents drug tolerance and drug resistance in *Mycobacterium tuberculosis*. Proc Natl Acad Sci U S A *114*, 4495-4500.

**Table S1: Strains used in this study.**

| **Strain Name** | **Source** |
|---|---|
| iBMDM | Gift of Dr. Christopher Sassetti |
| *M. smegmatis* mc$^2$155 | NC_008596 in GenBank |
| *M. smegmatis ΔsugC* | Gift of Dr. Rainer Kalscheuer |
| *M. smegmatis CΔsugC* | Gift of Dr. Rainer Kalscheuer |
| *M. smegmatis ΔotsA* | Gift of Dr. Rainer Kalscheuer |
| *M. smegmatis ΔtreYZ* | Gift of Dr. Rainer Kalscheuer |
| *M. smegmatis ΔtreS* | Gift of Dr. Rainer Kalscheuer |
| *M. smegmatis Δtre* | Gift of Dr. Rainer Kalscheuer |
| *M. smegmatis ΔotsA* pYAB-*tre* | This study |
| *M. smegmatis ΔtreYZ* pYAB-*tre* | This study |
| *M. smegmatis* pYAB | Gift of Dr. Yasu Morita |
| *M. smegmatis* pYAB-*tre* | This study |
| *M. tuberculosis* H37Rv | Gift of Dr. Rainer Kalscheuer |
| *M. tuberculosis ΔsugC* | Gift of Dr. Rainer Kalscheuer |
| *M. tuberculosis CΔsugC* | Gift of Dr. Rainer Kalscheuer |
| *E. coli* XL-1 blue | Agilent technologies |

### Figure S1

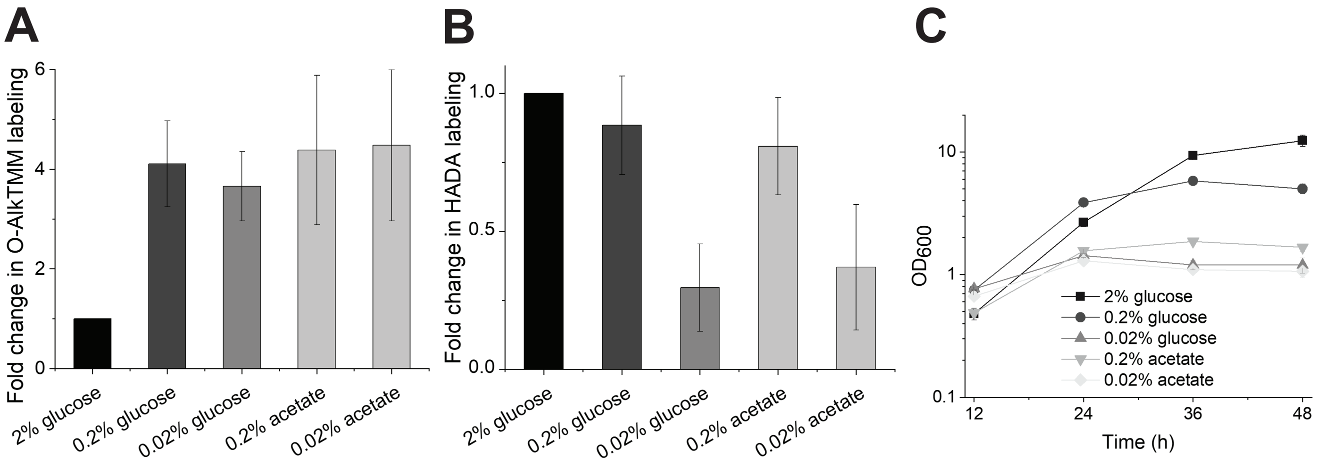
A
Fold change in O-AlkTMM labeling
0
2
4
6
2% glucose
0.2% glucose
0.02% glucose
0.2% acetate
0.02% acetate
B
Fold change in HADA labeling
0.0
0.5
1.0
2% glucose
0.2% glucose
0.02% glucose
0.2% acetate
0.02% acetate
C
OD600
0.1
1
10
12
24
36
48
Time (h)
2% glucose
0.2% glucose
0.02% glucose
0.2% acetate
0.02% acetate

### Figure S2

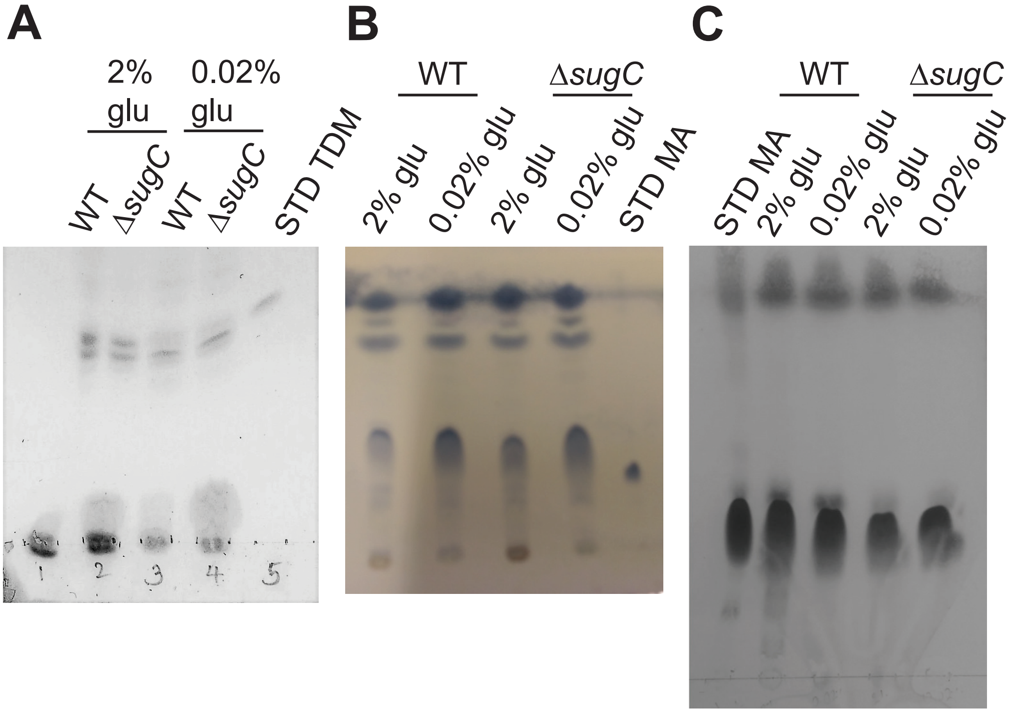
A
2% glu
0.02% glu
WT
ΔsugC
WT
ΔsugC
STD TDM
1
2
3
4
5
B
WT
ΔsugC
2% glu
0.02% glu
2% glu
0.02% glu
STD MA
C
WT
ΔsugC
STD MA
2% glu
0.02% glu
2% glu
0.02% glu

### Figure S3

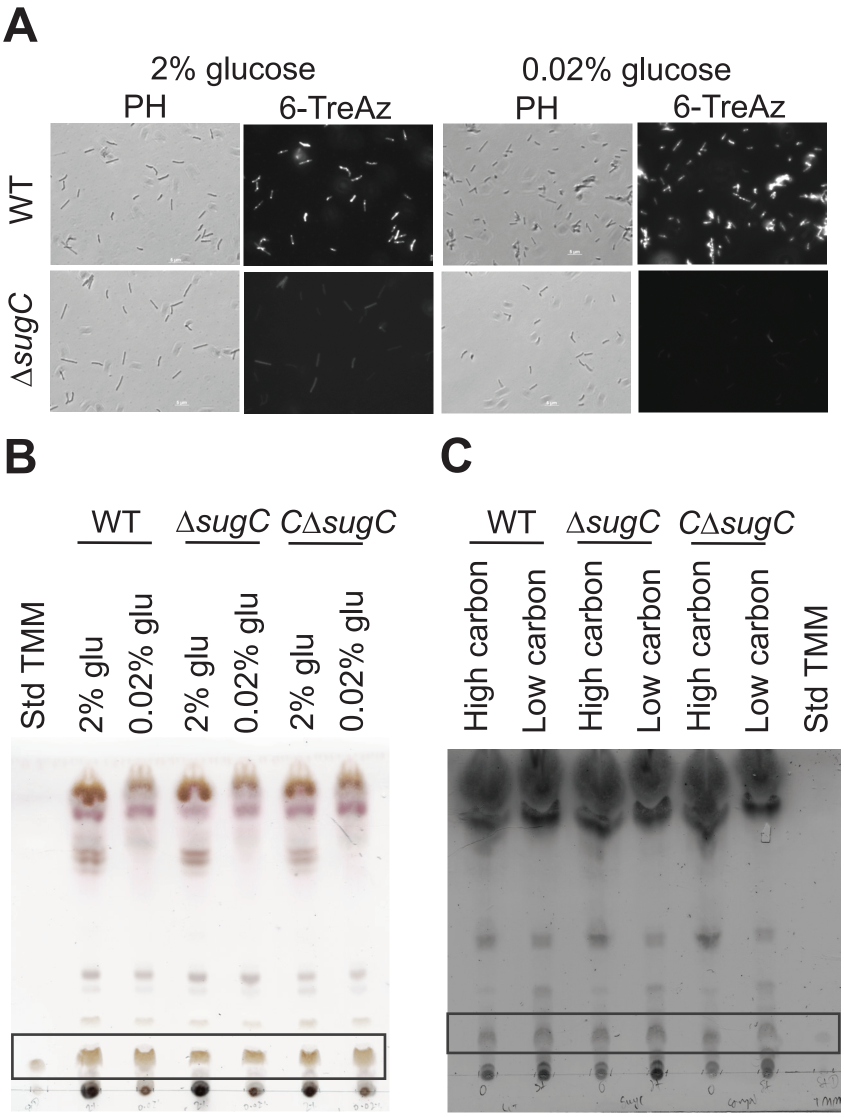
A
2% glucose
0.02% glucose
PH
6-TreAz
PH
6-TreAz
WT
ΔsugC
B
WT
ΔsugC
CΔsugC
Std TMM
2% glu
0.02% glu
2% glu
0.02% glu
2% glu
0.02% glu
C
WT
ΔsugC
CΔsugC
High carbon
Low carbon
High carbon
Low carbon
High carbon
Low carbon
Std TMM

### Figure S4

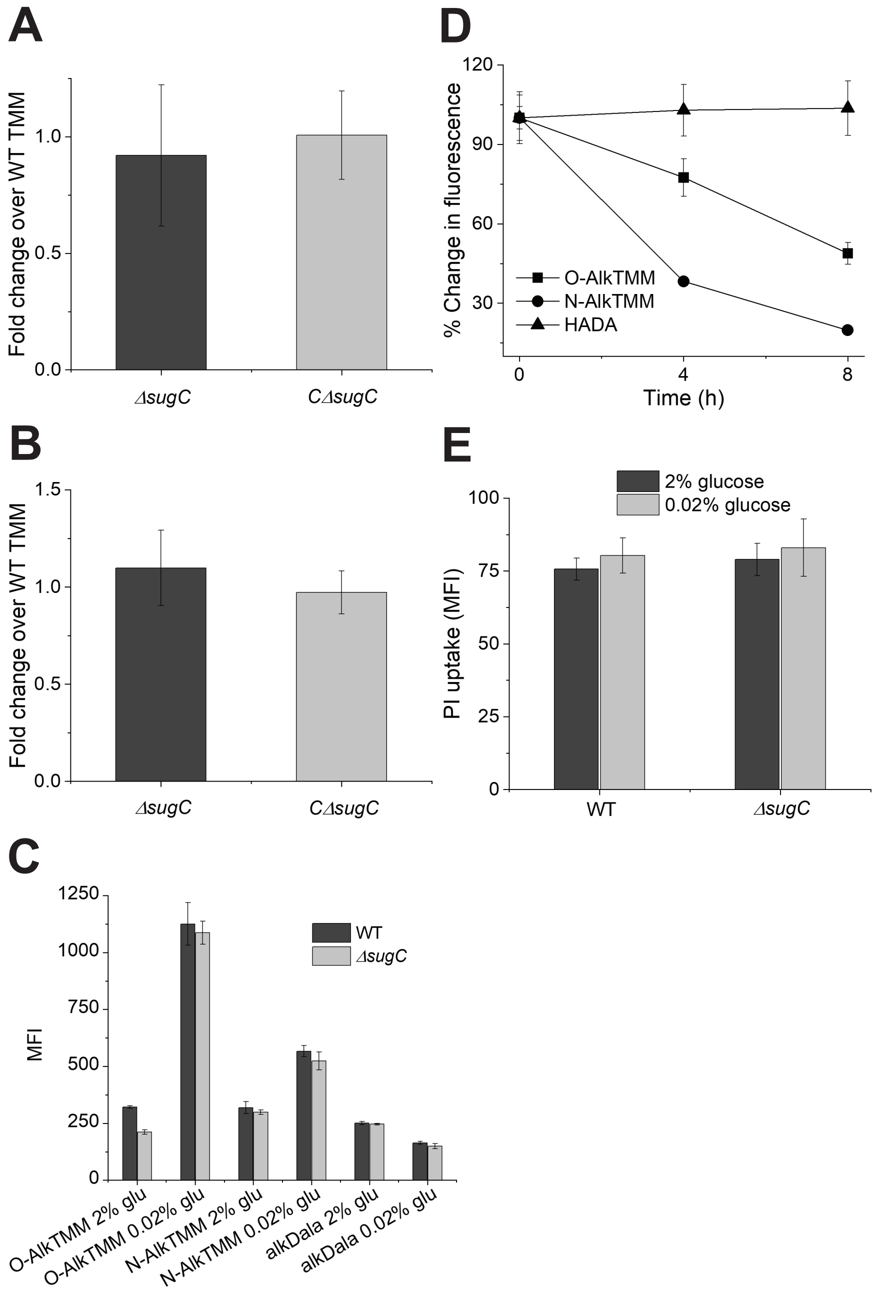

A
Fold change over WT TMM
ΔsugC
CΔsugC
D
% Change in fluorescence
Time (h)
O-AlkTMM
N-AlkTMM
HADA
B
Fold change over WT TMM
ΔsugC
CΔsugC
E
PI uptake (MFI)
2% glucose
0.02% glucose
WT
ΔsugC
C
MFI
WT
ΔsugC
O-AlkTMM 2% glu
O-AlkTMM 0.02% glu
N-AlkTMM 2% glu
N-AlkTMM 0.02% glu
alkDala 2% glu
alkDala 0.02% glu

### Figure S5

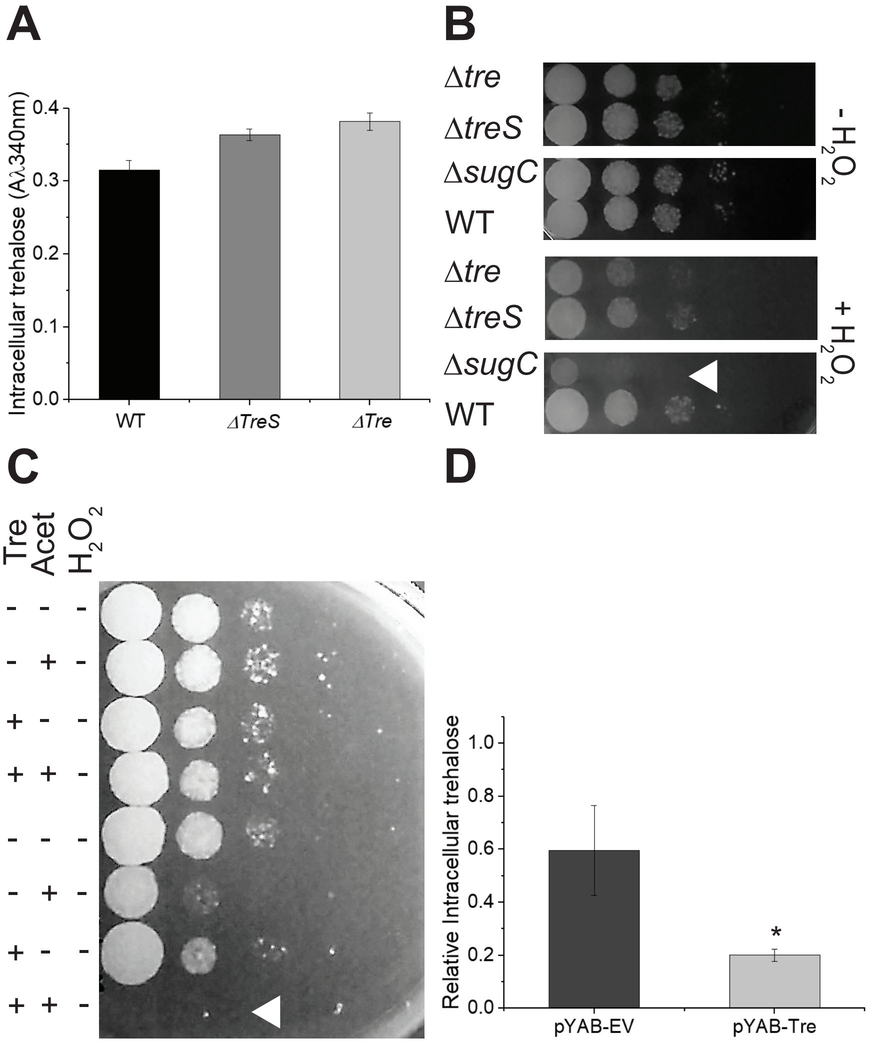
A
Intracellular trehalose (Aλ340nm)
0.0
0.1
0.2
0.3
0.4
WT
ΔTreS
ΔTre
B
Δtre
ΔtreS
ΔsugC
WT
Δtre
ΔtreS
ΔsugC
WT
- H2O2
+ H2O2
C
Tre
Acet
H2O2
- - -
- + -
+ - -
+ + -
- - -
- + -
+ - -
+ + -
D
Relative Intracellular trehalose
0.0
0.2
0.4
0.6
0.8
1.0
*
pYAB-EV
pYAB-Tre
